# *Plasmodiophora brassicae* in its environment-effects of temperature and light on resting spore survival in soil

**DOI:** 10.1101/819524

**Authors:** Kher Zahr, Alian Sarkes, Yalong Yang, Qixing Zhou, David Feindel, Michael W. Harding, Jie Feng

**Author notes:** Correspondence to: Michael W. Harding, Jie Feng.

## Abstract

Clubroot caused by *Plasmodiophora brassicae* is an important disease on cruciferous crops worldwide. Management of clubroot has been challenging, due largely to the millions of resting spores produced within an infected root that can survive dormant in the soil for many years. This study was conducted to investigate some of the environmental conditions that may affect the survival of resting spores in the soil. Soil samples containing clubroot resting spores (1 × 10^7^ spores g^-1^ soil) were stored at various temperatures for two years. Additionally, other samples were buried in soil, or kept on the soil surface in the field. The content of *P. brassicae* DNA and the numbers of viable spores in the samples were assessed by quantitative polymerase chain reaction (qPCR) and pathogenicity bioassays, respectively. The results indicated that 4°C, 20°C and being buried in the soil were better conditions for spore survival than were −20°C, 30°C and at the soil surface. Most of the spores kept on the soil surface were killed, suggesting the negative effect of light on spore viability. Additional experiments confirmed that ultraviolet (UV) light contributed a large negative effect on spore viability as lower pathogenicity and less DNA content were observed from the 2-and 3-hour UV light treated spores compared to the untreated control. Finally, this work demonstrated that DNA-based quantification methods such as qPCR can be poor predictors of *P. brassicae* disease potential due to the presence and persistence of DNA from dead spores.

## Introduction

Clubroot, caused by the protist *Plasmodiophora brassicae* Woronin, is an important threat to Canadian canola (*Brassica napus* L.) production [1]. In the Canadian Prairies, clubroot was first identified on canola in 2003 in a dozen fields near Edmonton, Alberta [2]. The disease has spread throughout much of the canola producing areas of Alberta [3,4], and has also been confirmed in canola fields in Saskatchewan [5], Manitoba [6], Ontario [7] and North Dakota [8].

Many strategies have been proposed for clubroot management and amongst those, crop rotation, along with the use of clubroot-resistant cultivars, was demonstrated to be effective [9]. However, in fields with high resting spore populations, the selection for virulent, but rare, *P. brassicae* pathotypes can lead to a shift in the population. The once rare virulent pathotype(s) becomes predominant and new sources of resistance must then be found. Shifts toward virulent pathotypes have been reported in Alberta since 2014 [10,11]. As a result of these findings, crop rotation (2-3 year break from a host crop), coupled with deployment of cultivars with genetic resistance, have been strongly recommended as the most reliable and sustainable clubroot management method [12,13].

The environmental effects on *P. brassicae* and clubroot have been well documented [14-18]. However, these reports have measured the effects of environmental and soil parameters on the germination, infection, severity of disease and root gall decay. Few studies have evaluated the effects of the soil environment on resting spore viability. The resting spores, which start the initial infection, have been reported to survive in the soil for up to 20 years, with a half-life of about 4 years [19]. More recently, there is evidence that, rather than a half-life, the resting spores experience a sharp decline in viability during the first two years without a host [12,20] followed by a slow decline of spore viability over the next 10 to 20 years. Regardless of the kinetics of spore degradation, little is known about which environmental parameters have the greatest effect(s) on spore survival. During the prolonged surviving time, the viability of spores may be impacted by soil type, soil pH values, water content, temperature and light [14]. Other factors may include the number of freeze-thaw cycles, rapid temperature shifts, microbial activity, population size, premature signals to exit dormancy, etc [14]. There is almost no information on how these soil environment parameters may affect the long-term viability of *P. brassicae* resting spores. In one study, the soil pH was found to play an important role in spore viability, and that soil temperature and moisture had much less effect [21]. However, the duration of the environmental exposure was only 30-days leaving us without an understanding of long-term effects of the soil environment on resting spore survival.

Since *P. brassicae* resting spores generally reside underground and are non-motile, most spores produced during one growing season remain in situ as dormant inoculum able to survive adverse environmental conditions or prolonged absences of host plants. Thus, in locations where resting spores have been present for many years, it is understood that while some spores survive, many others die beside the living ones. With this in mind, studies on clubroot diagnostics and clubroot management using the amount of DNA assessed by polymerase chain reaction (PCR) or quantitative PCR (qPCR) to represent the population of living spores in plant or soil samples could result in inaccurate conclusions. This inaccuracy results from the inability to ensure that only DNA from living cells is amplified by PCR, and thus the contribution of DNA from dead spores to quantification experiments is unknown. To address this complication, we evaluated the survival of resting spores using qPCR analysis side-by-side with pathogenicity bioassays. The results from the two approaches were compared and any inconsistency was further investigated. The objectives of this study were 1) to assess the viability of resting spores after being stored in different conditions for two years, 2) to identify any differential effects of environmental conditions on the survival of spores, 3) to illustrate the difference between DNA content and the biological potential (number of living spores) by using the same samples assessed by qPCR and pathogenicity bioassays, and 4) improve our understanding of *P. brassicae* survival, biomass assessments, and clubroot epidemiology.

## Materials and methods

### Ethics statement

No specific permission was required for the fields from which the *P. brassicae* populations were derived. All of the field studies were carried out in a closed and protected green house or a growth chamber in Crop Diversification Centre North (CDCN). This study did not involve endangered or protected species.

### *Plasmodiophora brassicae* populations

Two *P. brassicae* populations, one pathotype 3H and the other composed mainly of pathotype 5X, were used exclusively in this study. The pathotype 3H population was the Led09 strain [22] and was determined to be pathotype 3 by its reactions on the Williams’ deferential hosts [23], and pathotype H according to reactions on the Canadian Clubroot Differentials [24]. The pathotype 5X population was collected from a canola field in Sturgeon County, Alberta in 2016 and later was found to be pathotype 5X by using the specific PCR primers developed by Zhou et al. [25]. These two populations were maintained on the susceptible canola cultivar Westar in a greenhouse at CDCN. In the greenhouse, galls were collected from inoculated plants 42 days after inoculation. Galls were stored at −20°C for one month, and then chopped into 1 cm^3^ cubes, which were mixed at a ratio of 50% pathotype 3H to 50% pathotype 5X. This mixture was used to prepare resting spores for soil inoculation immediately or after being stored at −20°C for two years.

### Preparation of soil samples

A 2 kg black soil sample was collected from a research field plot at CDCN. The soil was autoclaved twice for one hour at 121°C, 15 psi and air dried at room temperature for three days. The dried sample was ground with a coffee grinder and then aliquoted into 1.5-mL microcentrifuge tubes with 1 g soil per tube. The tubes were stored at −20°C.

### Soil inoculation with resting spores

Resting spores were prepared from samples of the gall mixture according to Zhang et al. [26]. The spore suspension was adjusted to 2 × 10^8^ mL^-1^ and from which dilutions were prepared. For soil inoculation, 50 µL of a spore suspension was added into one of the 1.5-mL tubes containing 1 g soil, which ensured that all inoculated soil samples had the same water content (50 µL g^-1^ soil) despite different spore concentrations. After inoculation, the tubes were kept at room temperature for one hour to allow the inoculum to be absorbed evenly in the soil sample. The tubes were then vortexed for 10 seconds and immediately used in the subsequent experiments. For simplicity, these tubes will be referred to as ‘original-tubes’ throughout the rest of this paper.

### Treatments of resting spores in different environmental conditions

The original-tubes were prepared at the final concentration of 1 × 10^7^ spores g^-1^ soil. One hundred of the original-tubes were sealed in a 1-L glass media bottle (Fig. 1A), and each bottle was kept in one of the following conditions: 1) buried in soil at 10 cm beneath the soil surface in a field at CDCN, 2) on the surface of bare soil in the same field, 3) in an incubator set at 30°C, 4) in a growth chamber set at 20°C, 5) in a 4°C refrigerator and 6) in a −20°C freezer. All bottles except the one on soil surface were wrapped with double-layers of aluminum foil to ensure darkness and to protect the bottle. After approximately two years (23.5 months), tube samples were collected from each treatment and the soil in the tubes were subjected to qPCR analyses for measurement of *P. brassicae* DNA, and plant inoculation bioassays for measurement of pathogenicity.

**Fig 1.**
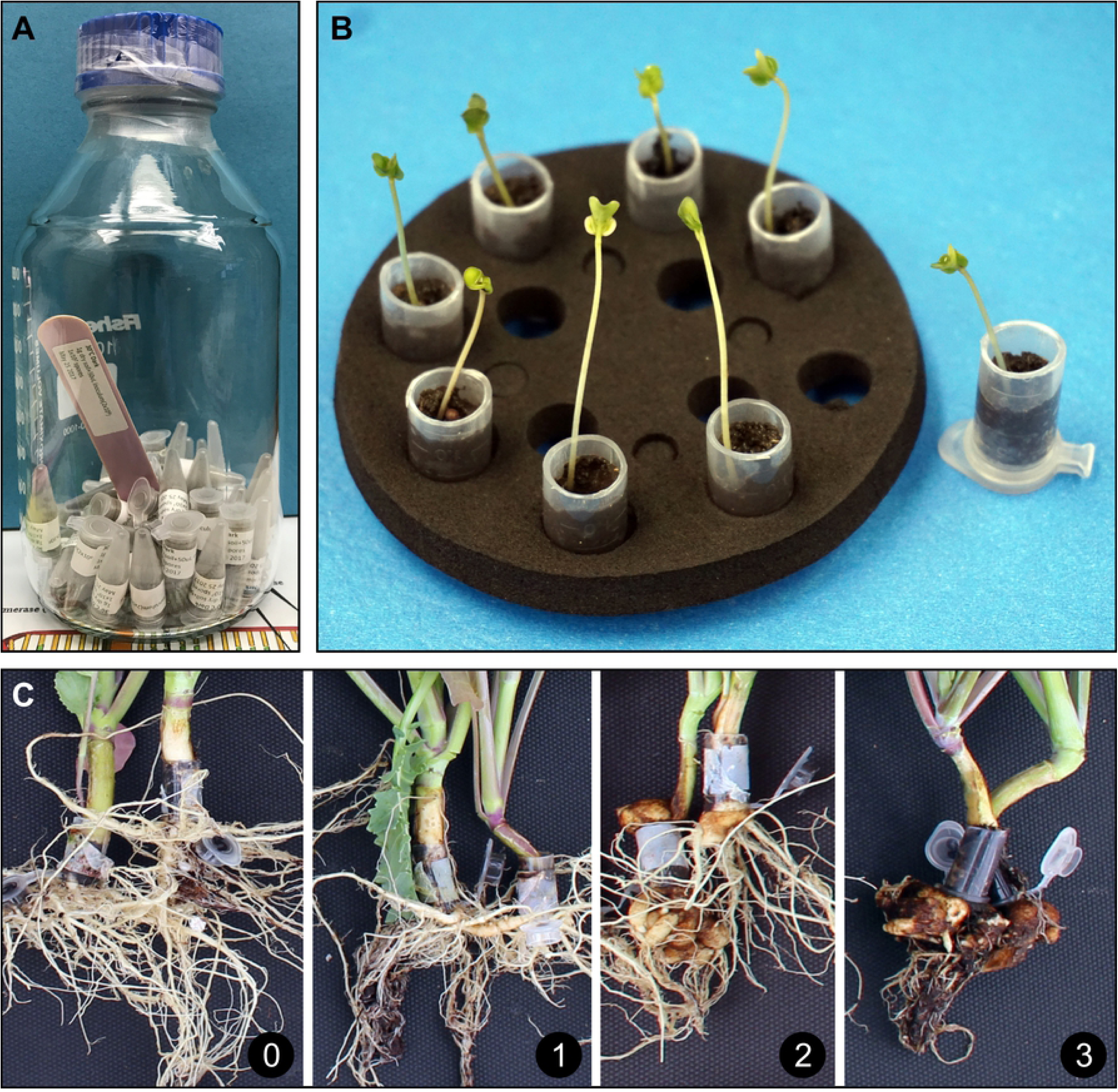
The tube-method for *Plasmodiophora brassicae* resting spore treatments and pathogenicity assay. Tubes each containing 1 g soil inoculated with 1 × 10^7^ resting spores in glass bottle ready for environmental treatment (A) and methods for pathogenicity testing of the treated soil samples (B and C). B, Canola seedlings inoculated with soil samples containing resting spores. C, the 0-to-3 clubroot rating scale.

### Autoclave treatment

Original-tubes were prepared at the final concentration of 1 × 10^7^ spores g^-1^ soil. With their caps open, the tubes were autoclaved with the dry cycle mode at 121°C, 15 psi for 0.5, 1, 2 or 3 hours. After autoclave, the tubes were kept at 4°C for no more than 24 hours before subjected to qPCR analyses.

### Ultraviolet light treatment

For UV light treatment, original-tubes were prepared at the final concentration of 1 × 10^6^ spores g^-1^ soil. Then the soil from each tube was transferred into a 5-cm petri dish. With the lid closed, the petri dish was shaken by hand for 30 seconds to make sure that the soil and the inoculum were well-mixed. After removing the lid, the petri dish was placed in a PCR workstation (Model P-048-02; CBS Scientific, San Diego, CA) and treated with UV-light for 1, 2 or 3 hours. The petri dishes were 15 cm beneath the 30-watt UV lamp, which resulted in a 400 µW/cm^2^ UV intensity being applied to the samples, as calculated according to the manual of the PCR workstation. After UV treatment, the petri dishes were kept under fluorescent light in an alternative biosafety hood until the complement of the 3-hour UV treatment. Then the soils of all treatments were transferred back to the 1.5-mL tubes and kept at 4°C for no more than 24 hours before being subjected to qPCR analyses and pathogenicity bioassays.

### DNA extraction from soil samples

Before sampling for DNA extraction, the 1-g soil sample in each 1.5-mL tube was homogenized by transferring the soil into a Ziploc bag and rubbing the bag by hand. From each bag 100 mg of soil was collected. Using the DNeasy PowerSoil kits (Qiagen Canada, Toronto, ON), genomic DNA was extracted from these 100-mg samples with a QIAcube instrument (Qiagen). The resultant DNA from each soil sample was eluted in 50 µL elution buffer included in the PowerSoil kit. DNA concentration was not measured.

### Quantitative PCR analysis

A pair of qPCR primers (CrqF2/CrqR2) was manually designed according to the rDNA sequence of a *P. brassicae* strain (GenBank accession number: KX011135). In this study, all PCR analyses were probe-based with the combination of this primer pair and the fluorescently labeled probe PB1 [27]. The sequences of CrqF2, CrqR2 and PB1 are: CTAGCGCTGCATCCCATATC, TGTTTCGGCTAGGATGGTTC and 6-FAM/CCATGTGAA/ZEN/CCGGTGAC/3IABkFQ, respectively. The primers and probe were synthesized by Integrated DNA Technologies (IDT; Coralville, IA). All qPCRs were conducted in PrimeTime Gene Expression Master Mix (IDT) using a CFX96 Touch Real-Time PCR Detection System (Bio-Rad Canada, Montreal, QC). Each 15 µL qPCR reaction contained 0.5 µM of each primer, 0.18 µM probe and 2 µL template DNA. The qPCR program consisted of 40 cycles of denaturation at 95°C for 10 sec (3 min for the initial denaturation) and annealing/extension at 60°C for 30 sec.

### Pathogenicity bioassays

The conical part of the 1.5-mL tubes containing 1 g inoculated soil was cut off. The tubes were sited upside-down on a floating tube rack (Fig. 1B) and 500 µL of water was added to each tube. Seedlings of canola cultivar Westar, generated from seeds on moist filter paper under continuous light for 96 hours, were transferred into the tubes with one seedling per tube. The tubes were incubated in a growth chamber maintained at 24°C/18°C (day/night) with a 16-h photoperiod. After five days, the cap of the tube was opened and the tube was transplanted into a 1.9-L square pot filled with Sunshine mix #4 potting soil (SunGro Horticulture, Agawam, MA). The pots were kept on individual plates to prevent cross-contamination in the greenhouse maintained at 24°C/18°C (day/night) with a 16-h photoperiod and watered from the bottom every second day with tap water at pH 6.4 (adjusted with HCl). After 42 days, the plants were separated into classes using a 0-to-3 scale [28], where 0 = no clubbing, 1 < one-third of the root with symptoms of clubbing, 2 = one-thirds to two-thirds clubbed, and 3 > two-thirds clubbed (Fig. 1C).

### Experimental design

The initiation of the resting spore storage treatments and the construction of the qPCR standard curve were conducted in May 2017. Other experiments were conducted in or after April 2019. All qPCR analyses and pathogenicity assays were conducted twice with similar results. Data from both experiments of the pathogenicity assay on the 2-year-stored spores were included. For other pathogenicity assays and all qPCR analyses, data from only one experiment was included. In each qPCR analysis, template DNA was extracted from three biological samples (three 100 mg soil samples from alternative tubes) for each treatment. For each DNA, qPCR was conducted with three technical repeats. The averaged data from these three technical repeats was treated as one data point in the statistical analysis. In each pathogenicity assay, each treatment was tested on five biological repeats (tubes). After transplanting, the pots in each experiment were kept on the greenhouse benches as randomized complete blocks.

### Statistics

Data from each experiment was subjected to analysis of variance (ANOVA) using Microsoft Excel. Based on the ANOVA results, differences between treatments were assessed with the Fisher’s LSD test (P ≤ 0.05) using the Excel add-in DSAASTAT developed by Dr. Onofri at the University of Perugia, Italy (http://www.casaonofri.it/repository/DSAASTAT.xls).

## Results

### Accuracy of the quantitative PCR assay

A qPCR standard curve was constructed based on the quantification cycle (Cq) values of DNA templates extracted from soil samples containing a 10-fold dilution series (from 1 × 10^7^ to 1 × 10^3^ spores g^-1^ soil) of *P. brassicae* resting spores (Fig. 2). The regression equation of the standard curve is Y = −3.0650X + 40.7583 with R^2^ = 0.9968. The calculated primer efficiency is 112%. For any soil sample, if the Cq value was obtained, its spore concentration or the equivalent of spore concentration could be calculated using the regression equation.

**Fig 2.**
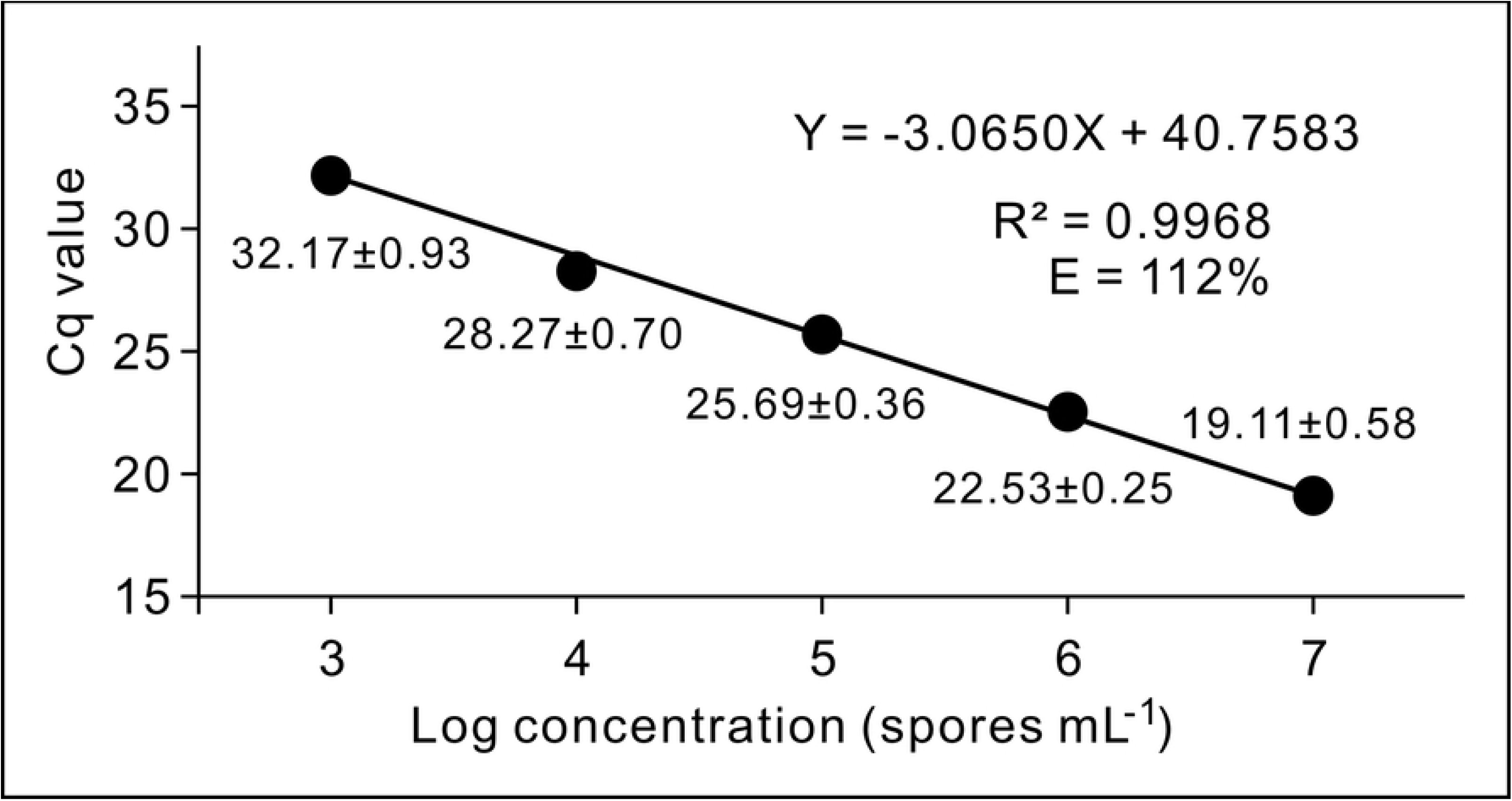
The standard curve generated from the mean of quantification cycle (Cq) values against log10 of spore concentrations in soil samples containing tenfold serial dilutions of *Plasmodiophora brassicae* resting spores. The regression equation, the R^2^ score of the equation and the efficiency of the primers (E), are indicated on the top of the curve. Each data point is shown as mean ± standard deviation (n=3).

### Quantitative PCR analysis of spores stored in different conditions for two years

After stored in different environmental conditions for two years, soil samples inoculated with *P. brassicae* resting spore were analysed by qPCR. Based on the Cq values, the equivalent spore concentration was calculated (Fig. 3). Compared to the fresh prepared 1 × 10^7^ spores g^-1^ soil control, soil samples stored at −20°C, 4°C or being buried in soil maintained a similar amount of *P. brassicae* DNA. In contrast, DNA in the soil samples stored at 20°C, 30°C or on soil surface was less than the control. In addition, DNA in the soil sample kept on soil surface was also less than the sample stored at −20°C or 4°C. These results indicated that 20°C, 30°C and the soil surface conditions are not favorable to the survival of *P. brassicae* resting spores and among them the soil surface conditions are most harmful. Furthermore, the DNA amount in a canola gall, before and after being stored at −20°C for two years was not different. When the mean Cq of the fresh prepared 1 × 10^7^ spores g^-1^ soil control (M = 19.48, SD = 0.69; Fig. 2) was compared to that of the soil sample prepared and assessed two years ago (M = 19.11, SD = 0.58; Fig. 1) by a Student’s t-test, no significant difference was found (t = 0.7109, p = 0.5164).

**Fig 3.**
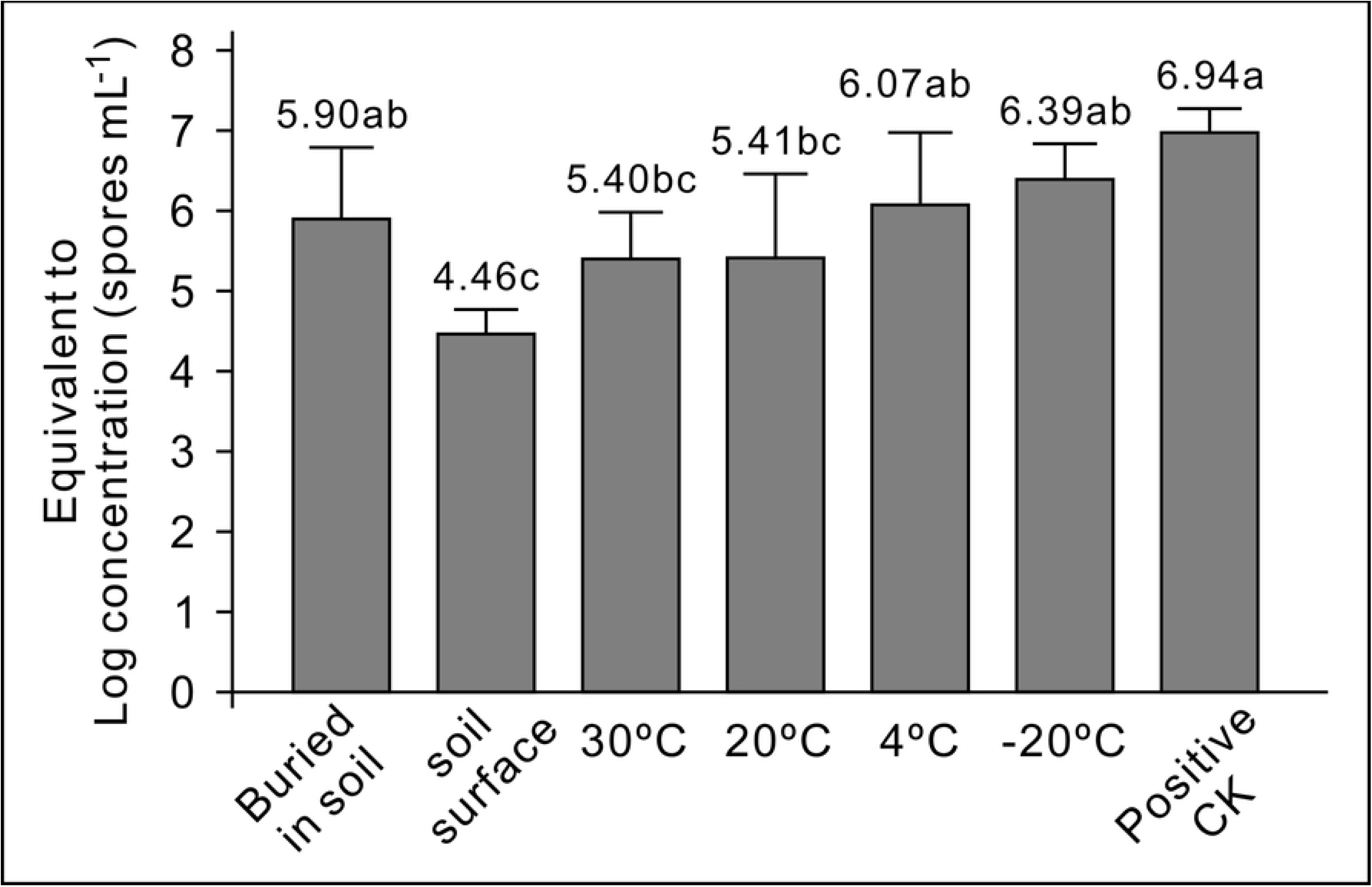
qPCR analyses of soil samples containing 1 × 10^7^ *Plasmodiophora brassicae* resting spores g^-1^ soil that have been stored in different conditions for two years. Using the mean Cq value of each sample obtained from qPCR and the regression equation in Fig. 2, the value of Y was calculated for each treatment and shown as the equivalent to log concentration. Means in the plot topped by the same letter do not differ based on Fisher’s LSD test at P ≤ 0.05 (n = 3).

### Pathogenicity assay of spores stored in different conditions for two years

The pathogenicity of spores in the soil samples was evaluated based on the clubroot severity on the susceptible canola cultivar Westar (Fig. 4A). In the two pathogenicity experiments, soil samples stored at 20°C, 4°C and being buried in soil showed similar pathogenicity to the fresh prepared 1 × 10^7^ spores g^-1^ soil control, while the remaining treatments showed a reduction in pathogenicity. The soil sample stored at 4°C showed a similar pathogenicity in one experiment and lower pathogenicity in the other experiment, compared to the fresh prepared 1 × 10^7^ spores g^-1^ soil control. No clubroot symptoms were observed from the soil sample kept on soil surface in both experiments. Interestingly, the soil sample stored at −20°C showed pathogenicity lower than the samples stored at 4°C, 20°C or buried in the soil.

**Fig 4.**
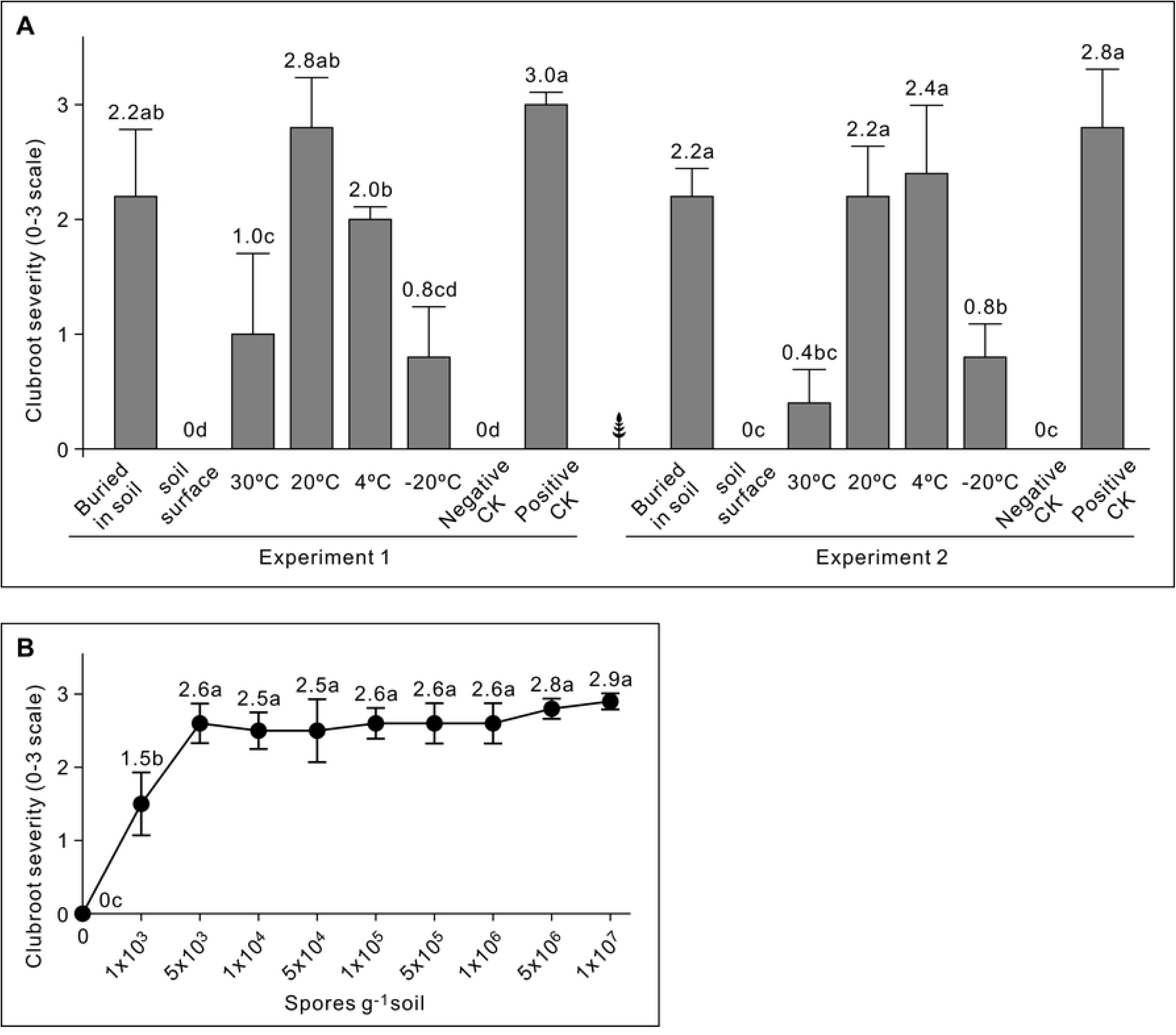
Pathogenicity bioassays of soil samples inoculated with *Plasmodiophora brassicae* resting spores. Means in the plot topped by the same letter do not differ based on Fisher’s LSD test at P ≤ 0.05 (n = 5). A, Two experiments on soil samples containing 1 × 10^7^ spores g^-1^ soil spores that had been stored in different conditions for two years. Negative CK, an uninoculated soil sample that has been stored at −20°C for two years; Positive CK, a fresh prepared soil sample containing 1 × 10^7^ spores g^-1^ soil spores. B, one experiment on fresh prepared soil samples containing a 5-fold dilution series (1 × 10^7^ to 1 × 10^3^ spores g^-1^ soil) of spores.

Difference in pathogenicity among the soil samples can be attributed to the fact that the numbers of viable spores in these samples were different. To estimate the disease potential of viable spores in these soil samples, pathogenicity bioassays on soil samples containing a 5-fold dilution series (1 × 10^7^ to 1 × 10^3^ spores g^-1^ soil) of resting spores were conducted in parallel with the pathogenicity assays of the stored samples. The results showed that a spore concentration ≥ 5 × 10^3^ spores g^-1^ soil could cause clubroot with same severity as the spore concentration at 1 × 10^7^ spores g^-1^ soil (Fig. 4B). Thus, we could conclude that the concentration of living spores in the soil samples kept on soil surface, at 30°C or at −20°C were lower than 5 × 10^3^ spores g^-1^ soil while the concentration in the sample on soil surface was lower than 1 × 10^3^ spores g^-1^ soil.

### Pathogenicity assays vs. quantitative PCR

Comparing the results from the qPCR analysis to those of the pathogenicity bioassays indicated that qPCR should not be used to assess the numbers of living spores or disease potential. When spores die, their DNA can remain in the sample and produce qPCR amplicons, although more or less of this DNA would degrade depending on the environmental conditions. This is illustrated by the qPCR and pathogenicity bioassay results of the soil sample stored at −20°C. The qPCR data indicated that the amount of DNA in the −20°C sample was equivalent to 1 × 10^6.39^ spores g^-1^ soil, which is the highest mean among all treatments except the fresh prepared 1 × 10^7^ spores g^-1^ soil control (Fig. 3). However, pathogenicity bioassays indicated that less than 5 × 10^3^ living spores g^-1^ soil were present in this sample. The difference between the qPCR result and the pathogenicity bioassay result of this sample was greater than those of the samples on the soil surface, or at 20°C or 30°C. This could be explained by the assumption that most spores in samples incubated at −20°C were dead, but −20°C provided an environment that better maintained the integrity of DNA released from the dead cells. To test the contribution of dead spores to the DNA content in the soil sample, an autoclave experiment was conducted in which soil samples containing 1 × 10^7^ spores g^-1^ soil were autoclaved and then analyzed by qPCR. After autoclaved for 0.5 or 1 hour, *P. brassicae* DNA could still be detected (Fig. 5) with mean Cq values at 31.28 ± 0.55 and 32.84 ± 0.60, respectively. After autoclaved for more than 2 hours, the mean Cq values of the samples reached to 36 and some technical repeats failed to generate a Cq value. This result indicated that autoclaving for 2 hours is required to destroy all *P. brassicae* DNA released from dead spores. The 2-hour autoclaving may not be necessary for preparing negative control material for pathogenicity bioassays because the widely used 1-hour autoclave is sufficient to kill all the resting spores. However, in experiments with PCR or qPCR being used to quantify spore numbers based on DNA, autoclaving the sample for more than 2 hours is recommended.

**Fig 5.**
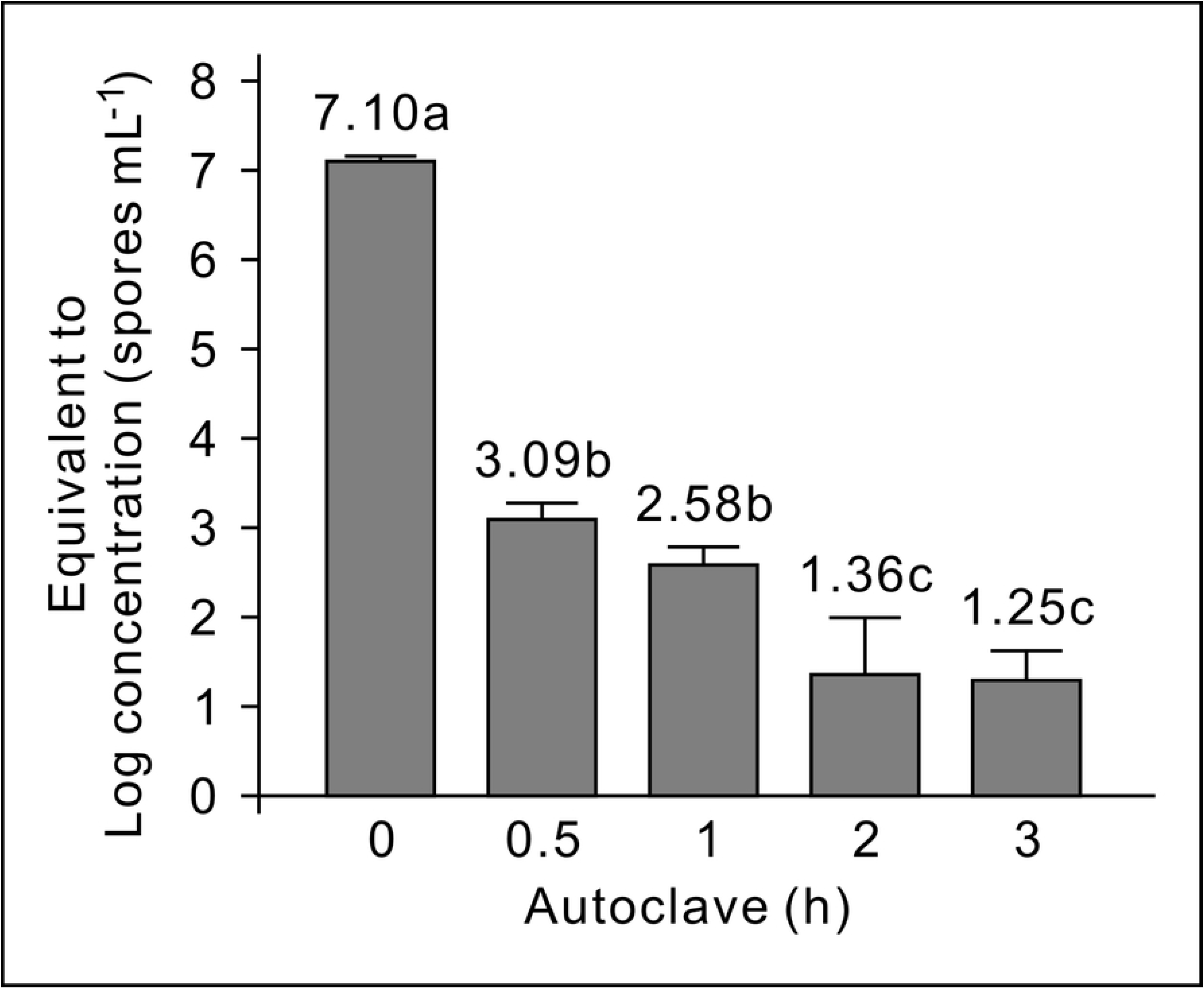
qPCR analysis of soil samples containing 1 × 10^7^ *Plasmodiophora brassicae* resting spores g^-1^ soil that have been autoclaved for 0, 0.5, 1, 2 or 3 hours. Using the mean Cq value of each sample obtained from qPCR and the regression equation in Fig. 2, the value of Y was calculated for each treatment and shown as the equivalent to log concentration. Means in the plot topped by the same letter do not differ based on Fisher’s LSD test at P ≤ 0.05 (n = 3).

### Ultraviolet light treatment

The pathogenicity difference between the sample buried in soil and that kept on the soil surface indicated that radiant sunlight may be able to kill resting spores. UV light, as a component of sunlight, is known to be harmful to life forms due to its ability to damage DNA [29]. To test the effect of UV light on the survival of the resting spores, soil samples containing 1 × 10^6^ spores g^-1^ soil were treated with UV light and subjected to qPCR analyses and pathogenicity bioassay. After treatment with UV light for one or two hours, a decrease in the quantity of DNA was observed (Fig. 6A). When the treatment time was extended to 3 hours, only 50% (10^5.66-5.94^) of DNA, compared to the non-treated control, could be identified from the soil sample, which indicated that more than half of the resting spores were killed by the 3-h UV light treatment. This result was supported by the pathogenicity bioassays (Fig. 6B), in which 2-and 3-hour UV light treated samples showed significantly reduced pathogenicity compared to the non-treated control.

**Fig 6.**
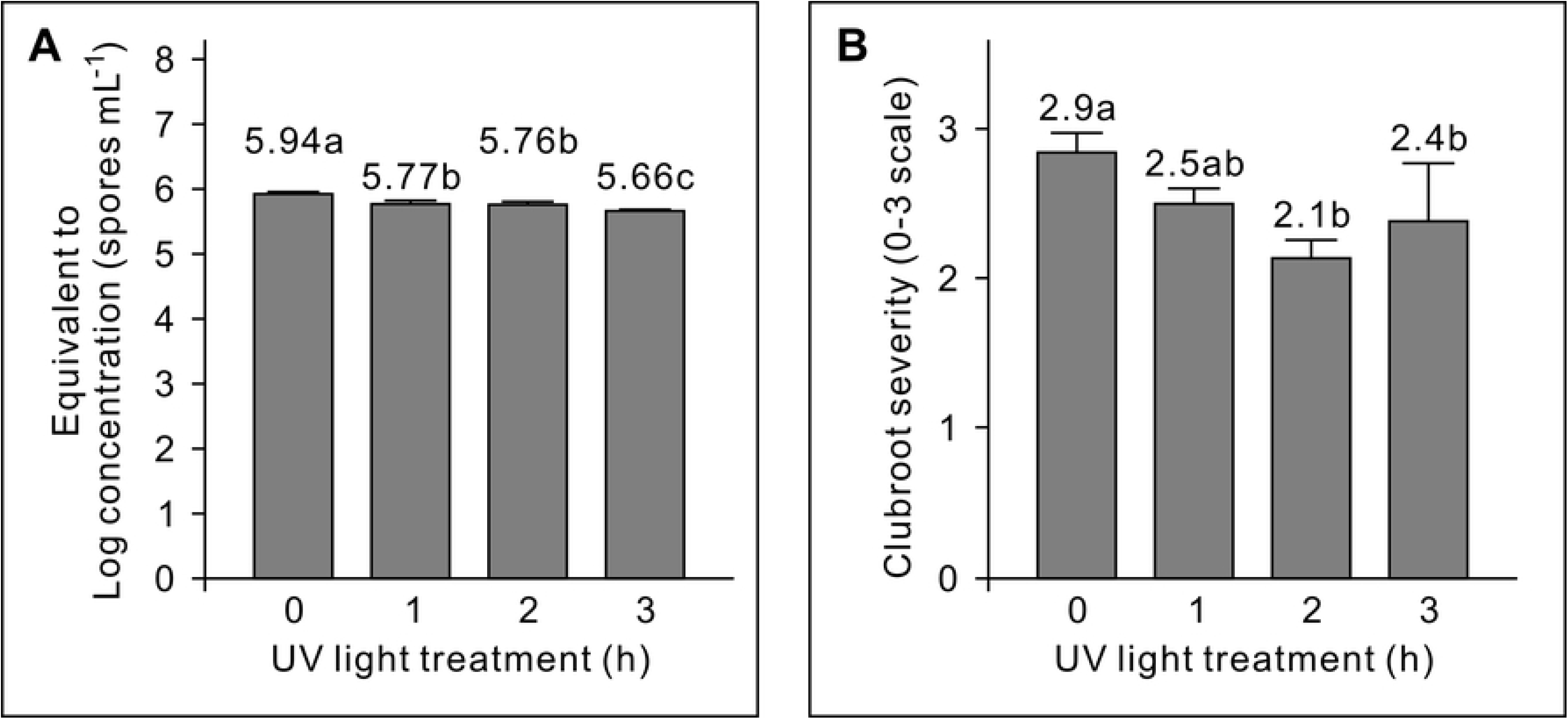
qPCR analysis (A) and pathogenicity assay (B) of soil samples containing 1 × 10^6^ *Plasmodiophora brassicae* resting spores g^-1^ soil that have been treated by UV light for 0, 1, 2 or 3 hours. Means in the plot topped by the same letter do not differ based on Fisher’s LSD test at P ≤ 0.05 (n = 3 for qPCR; n = 5 for pathogenicity assay). In the qPCR analysis, using the mean Cq value of each sample obtained from qPCR and the regression equation in Fig. 2, the value of Y was calculated for each treatment and shown as the equivalent to log concentration.

## Discussion

### Assessment of *Plasmodiophora brassicae* with quantitative PCR

In this study we used primer pair CrqF2/CrqR2 and probe PB1 for qPCR assessment of *P. brassicae* spores. The probe has been used with the primer pair DC1F/DC1mR [27,30], which was derived from Rennie et al. [31] but with the reverse primer modified. We redesigned the primers to avoid potential primer dimers as predicted by the Multiple Primer Analyzer (https://www.thermofisher.com). Compared to other published primers, this primer pair showed similar specificity and sensitivity to *P. brassicae* when used in SYBR green based qPCR (Feng, unpublished data). Since the sequence of the probe PB1 doesn’t create more specificity, we believed that this primer pair would be as efficient as others when used in probe-based qPCR.

The efficiency of the primers used for the template DNA in this study was 112%. This could be attributed to the PCR inhibitors present in the DNA template solution. For example, if the standard curve was created from a dilution series of one genomic DNA preparation with inhibitors (generally chemicals used for DNA extraction), primer efficiency larger than 100% would be expected, due to the fact that the inhibitors in the original template solution were diluted along with DNA dilution. However, in this study, the standard curve was created by serial DNA preparations from soil samples inoculated with spore dilutions. Thus, we concluded that at least some of the PCR inhibitors were derived from *P. brassicae* spores. The more spores from which the DNA derived, the more inhibitors in the DNA solution.

It has not escaped our notice that the standard deviations of qPCR data from each treatment in the UV light experiment (Fig. 6A) were smaller than those in the 2-year-storage experiments (Fig. 3). We inferred that in the 2-year-storage experiments, stochastic factors had more time to interact with the spore samples. Therefore, more variability was created within the different tubes, although these tubes were kept in the same bottle.

In this study, we provided the evidence that using qPCR data to enumerate or estimate living spores can, in some cases, dramatically overestimate the numbers of viable resting spores and disease potential. Few studies have been conducted to evaluate differences between using qPCR results and pathogenicity bioassay results to assess *P. brassicae* resting spore viability. PMA-PCR has been used for the assessment of *P. brassicae* living spores [32,33], however while this method has the potential to improve the measurement of viable spores via qPCR, the usefulness of PMA-PCR on quantification of absolute spore numbers and the standardization of the procedure, especially the step of photoactivation, still need to be improved. Additionally, the effects of photoactivation on living spores may have deleterious consequences as shown in this study with UV light experiments. Unfortunately, the results of the experiments reported herein suggest that the best way to measure spore viability and disease potential is pathogenicity bioassays, with vital staining of spores being a next best option [33].

### Survival of *Plasmodiophora brassicae* spores in different temperatures

Among the four controlled temperatures, 4°C and 20°C were better than −20°C and 30°C on maintaining spore viability (Fig. 4A). These results indicated that compared to 4°C and 20°C, continuous freezing, or high temperature (>20°C) are more harmful to the spores. It has been suggested that spores might be stored as dense suspensions at 3-4°C for 3 years without loss of viability [34], which was supported by the results of the soil sample kept at 4°C. In contrast, it has been a standard practice to store galls at −20°C for several years as stock inoculum [35], which was not supported by the results of the soil sample kept at −20°C. Furthermore, in this study, the spores buried in soil, which would have been continuously frozen during the two winter seasons, still maintained full viability. We attributed the disagreement between our results and those previously reported to the different water content in the samples. All our soil samples contained 5% water before being stored in different conditions. Samples in the −20°C treatment were frozen as soon as the treatment was started. In contrast, samples in the field treatment were not frozen for several months until the winter came, which likely allowed the soil samples to lose most of their water content. The tubes were sealed in bottles, but not with an air-tight seal. Frozen as plant galls or in dry samples may minimize the occurrence of cell crystallization thus more cells will survive. The samples buried in the soil might also experience high temperatures, but it should have been only intermittently with short time periods in the two Edmonton summer seasons.

### Survival of *Plasmodiophora brassicae* spores in field conditions

In many plant pathogens, a compensation for being soil-borne is to produce larger numbers of thick-walled, long-lived spores. The resting spores of *P. brassicae* have evolved to retain viability in the soil environment despite exposure to many seasons of adverse weather, and temperature/moisture extremes. Field studies indicated that *P. brassicae* resting spores have a half-life of at approximately 3.6 years and viable spores might exist for 20 years in the absence of suitable hosts before spore populations were eroded to undetectable levels [19]. More recently some experiments have suggested that the decay may not be linear, but rather a rapid decline in viability occurs over the first year or two without a host, followed by a much slower decline over the next 10 to 20 years [12,20]. In this study, we found that most spores can maintain full viability and pathogenicity after being buried in the soil for two years. In contrast, most spores at the soil surface totally lost their viability and pathogenicity after two years. However, since spores were kept in 1.5-mL tubes and the tubes were kept in bottles, results from either treatment cannot be used to predict the fate of spores in real field conditions. Nevertheless, the differences between the two treatments provided evidence that spores buried in the soil had more chances to survive than those on soil surface. According to Kim et al. [36], more than 97% of the total of *P. brassicae* inoculum was present in the surface soil (0-5 cm depth) and few resting spores were found below 40 cm. Cranmer et al. [37] indicated that genomic DNA of *P. brassicae* equivalent to 10^5^ spores g^-1^ soil of resting spores could be detected below the plow layer. Thus, in field conditions, pathogenic populations are influenced by the spore concentrations on the soil surface and in the soil, with potentially drastic differences in their viability decline curves for the two locations. In this study, spores buried in the soil remained pathogenic for more than 2 years but those on the soil surface did not. Furthermore, the low viability of spores on soil surface suggested that tillage between rotated crops might actually improve clubroot management by exposing more spores to the damaging radiant sunlight present at the soil surface.

### Effect of ultraviolet light on spore viability

It has been known for many years that UV light has various effects on fungi and other microorganisms [38]. When DNA is exposed to UV radiation adjacent thymine bases will be induced to form cyclobutane pyrimidine dimers by the condensation of two ethylene groups at C-5 and C-6. Additionally, adjacent thymines can be linked between the C-4 residue and the C-6 of its neighbor. In either case, a “kink” is introduced into the DNA. Therefore, by exposing microorganisms to UV radiation, their DNA will be photo-damaged and will not be amplified by DNA polymerase [39]. As another consequence, the DNA can’t replicate and thus the cells die.

*Plasmodiophora brassicae* has developed the ability to form resting spores with mechanisms to resist adverse thermal and aqueous conditions. Among many other factors, melanin may play a role in resting spores for UV light resistance. Melanin is a broad term for a group of natural pigments and localized in cell walls in most organisms [40,41]. In *P. brassicae*, the L-Dopachrome biosynthesis pathway, which is involved in melanin synthesis, has been predicted by a bioinformatic study [42]. However, in natural conditions, especially outside agriculture, most resting spores of *P. brassicae* would be buried in the soil for their entire life. Thus, it is likely that *P. brassicae* did not evolve a strongly expressed UV light resistance during evolution. This was supported by the current study, in which the *P. brassicae* resisting spores were found to be sensitive to UV light treatment.

To prevent the spread of clubroot, sanitation of field equipment is important. Currently bleach is recommended as one of the most effective chemicals for inactivation of clubroot resting spores (https://www.canolawatch.org/2018/06/27/clubroot-disinfectants-bleach-is-best/). However, bleach is corrosive to metals, causes rubber to harden, stains and damages clothing and footwear. Considering the effect of UV light on the viability of resting spores as demonstrated in the current study, UV light treatment may be an alternative for equipment sanitation. Compared to other treatments, UV light treatment has several advantages. Firstly the cost will be low because no consumable materials are required; secondly, it will be easy to perform because no attention or action is required during the treatment process; and finally, it is environment-friendly. On the other hand, shortcomings include that UV light does not readily penetrate surfaces but functions on direct surfaces only. Furthermore, it may require long exposure times for good efficacy. Thus, we would leave this topic open and expect future applied or adaptive studies to take this further.

The observed damage by UV light also challenges the notion that the spread of *P. brassicae* can be due to resting spores moving with wind-blown dust and soil erosion [43,44]. The results presented herein on the damaging effects of exposure to sunlight or UV would suggest that most spores moving with wind-blown soil occur at the soil surface, and likely inactivated after prolonged exposures to sunlight.

### Inoculating canola with *Plasmodiophora brassicae*-infected soil

As with other biotrophic plant pathogens, plant inoculation with *P. brassicae* can be challenging. In diagnosis service activities and studies on the pathogenicity of multiple strains (e.g. pathotyping, screening for resistant germplasms), inoculum sources are generally limited by the amount of *P. brassicae*-containing soil or root galls. Using small amounts of inoculum allows for more experimental replications to be set up, but often decreases the efficiency of infection. In this study, we developed and optimized a simple procedure for canola seedling inoculation (Fig. 1B and C; termed tube-method), which is extremely useful when the inoculum source is limited. Compared to the soil inoculation method [45], the tube-method uses much less inoculum; compared to the sealing-dipping method [46], the tube-method has higher levels of consistency among repeats and efficiency of infection. We recommend this inoculation method, regardless of whether the original inoculum is soil or root gall, in *P. brassicae* pathogenicity assays conducted in either labs or greenhouses.

## Author Contributions

Conceptualization: Jie Feng, Michael W. Harding.

Data Curation: Jie Feng.

Formal Analysis: Jie Feng.

Funding Acquisition: Jie Feng, Michael W. Harding, David Feindel.

Investigation: Kher Zahr, Alian Sarkes, Yalong Yang, Jie Feng, Qixing Zhou.

Methodology: Jie Feng, Yalong Yang.

Project Administration: Jie Feng.

Resources: Jie Feng, David Feindel.

Supervision: Jie Feng.

Validation: Jie Feng, Michael W. Harding.

Visualization: Jie Feng.

Writing – Original Draft Preparation: Jie Feng.

Writing – Review & Editing: Jie Feng, Michael W. Harding.

